# Hemoglobin C is prone to oxidative denaturation, resulting in red blood cell membrane damage in HbSC disease

**DOI:** 10.64898/2026.06.05.729662

**Authors:** Tahereh Setayesh, Anifat Tijani, Harsimran Kaur, Shristi Khanal, Zhenqi Zhu, Zachery Oestreicher, Katie Seu, József Balla, Mengna Chi, Russell E. Ware, Punam Malik

## Abstract

Sickle-hemoglobin-C (HbSC) sickle cell disease is characterized by RBC dehydration (xerocytosis), which promotes polymerization of HbS. HbSC causes substantial morbidity despite lower sickling potential than HbSS, suggesting a critical detrimental role of HbC in the disease pathophysiology. We derived HbCC mice by interbreeding our HbSC mice, which demonstrated a similar RBC phenotype of xerocytosis as humans with HbCC. We compared RBCs from HbCC, HbSC, and HbSS mice. Oxidized ferryl (Fe^4^+)-Hb, and its oxidative-denaturation, which results in hemichrome formation (Heinz-bodies), was most pronounced in HbCC>HbSC>HbSS, despite significantly higher reactive oxygen species in HbSS, illustrating a higher propensity of HbC to denaturation than HbS. RBC deformability followed a similar pattern, with Elongation Index lowest in HbCC<HbSC<HbSS. Next, we determined if RBC from HbSC patients on hydroxyurea showed improved membrane damage. Hydroxyurea treatment reduced Heinz-body formation and improved RBC deformability, despite negligible/modest fetal hemoglobin (HbF) induction, compared to non-hydroxyurea HbSC controls. The antioxidant quercetin showed a similar reduction in Heinz-body burden and improvement in RBC deformability as hydroxyurea, without affecting Hb or HbF concentration, reticulocyte count, or RBC xerocytosis. HbC-driven oxidative denaturation and membrane damage represent important contributors of RBC dysfunction in HbSC disease; hence, oxidative membrane injury could be targeted besides antisickling approaches.

## Introduction

Hemoglobin-Sickle hemoglobin-C (HbSC) is the second most common form of sickle cell disease (SCD) worldwide, affecting 30% of SCD patients. Although historically considered milder than the homozygous form, HbSS-SCD, HbSC is now being recognized to cause significant morbidity and mortality.^1-3^Additionally, HbSC-SCD results in higher incidence of proliferative retinopathy, thrombotic complications, and blood hyperviscosity than Hbss.^2-4^ HbSC (and HbCC) RBC are distinctly characterized by RBC dehydration (xerocytosis).^5-7^ Current dogma is that cellular dehydration results in increased HbS concentration within RBC, which promotes HbS polymerization even when HbS is at -50%/”trait” concentrations. However, polymerization is generally less extensive in HbSC than in HbSS RBC,^7,8^ suggesting additional deleterious effects of HbC.

Clinically, hydroxyurea reduces vaso-occlusive events in HbSC-SCD despite modest HbF induction.^9,10^ We recently showed that hydroxyurea improves HbSC RBC rheology in humanized HbSC mice, independent of fetal hemoglobin (HbF) induction, which was associated with reduced oxidized hemoglobin.^11^ These data suggested that oxidative damage to RBC may be an important and largely unrecognized contributor to HbSC pathophysiology.

Herein, using humanized HbSC, HbCC, and HbSS mice and blood samples from HbSC patients, we sought to determine whether HbC drives oxidative membrane damage. We show HbC has a higher propensity to denature than HbS, causing membrane damage, which can be mitigated both with hydroxyurea or antioxidant treatment.

## Methods

### Mice

Humanized Townes HbAA and HbSS mice, obtained from Dr. Timothy Townes, University of Alabama, and HbSC *(Hbb^tm2(HBG1,HBB*)TDW^/Hbb^eml(HBG1,HBB*)Malik^Hba^tml(HBA)TDW^/MalikJ)* **and HbCC** mice, generated in our lab,^11^, were studied under Institutional Animal Care and Use Committee— approved protocols at Cincinnati Children’s Hospital Medical Center. Mice (10-12 weeks-old) received hydroxyurea (50 mg/kg BW), quercetin (0.2 mg/kg BW), or the combination by intraperitoneal injection thrice weekly for 8 weeks before analyses. Mice (10-12 weeks old) received hydroxyurea (50 mg/kg BW), quercetin (0.2 mg/kg BW), the combination, or PBS vehicle control by intraperitoneal injection three times weekly for 8 weeks before analysis.

Human peripheral blood (PB) samples were obtained from patients with HbSC, HbCC, or HbC-13° thalassemia using Institutional Review Board-approved protocols.

### RBC studies

Hemoglobin denaturation and hemichrome formation were assessed by Heinz-body staining of PB. Heinz body-positive RBC were quantified, and ferryl Hb immune fluorescent staining was performed as previously described.^11-12^ RBC morphology was examined by scanning electron microscopy.^11^ RBC deformability was measured by Lorrca®-MaxSis ektacytometry using shear-stress, oxygen-gradient, and osmotic-gradient ektacytometry. Standard hematological indices, including hemoglobin, reticulocytes, and F-cell percentages, were measured as previously described.^11^

Statistical comparisons were performed using Mann-Whitney U test or by Analysis of Variance. Data are presented as mean± SEM. *P* values <0.05 were considered significant.

## Results and Discussion

We and Zhai et al. reported the generation of HbSC mice via gene editing of the Townes HbSS mice.^11-13^ We generated HbCC mice via interbreeding our HbSC mice. HbCC mice showed hematological characteristics typical of HbCC patients (see Supplementary Figure 1). RBCs and reticulocytes from HbCC mice were more severely dehydrated than HbSC>HbSS>HbAA RBCs, both by CHCM measurements and osmoscan ektacytometry (Supplementary Figure 1G-H, J-K). Scanning electron microscopy showed distinct RBC morphology across the different genotypes: HbAA RBC showed the expected biconcave discocytes, whereas HbSS, HbSC, and HbCC RBC exhibited progressively greater membrane and shape irregularities that occur with increasing xerocytosis (Figure 1A). HbC-containing RBC (HbSC and HbCC) frequently displayed distorted membranes and irregular surface contours. Additionally, shear-stress ektacytometry showed that RBC from both HbCC mice and humans (HbCC or HbCβ^0^-thalassemia) had the lowest deformability, compared to HbSC and HbAA RBC (Supplementary Figure 1L-M). The derivation of HbCC and HbSC mice from HbSS mice provides a unique, precisely controlled opportunity to compare mice that only differ by a single nucleotide at *HBB* codon 6, and to study the contribution of HbC to RBC pathobiology with or without the presence of HbS.

Oxidatively denatured hemoglobin results in hemichromes, which aggregate with the Band3 membrane protein and deposit on RBC membranes as Heinz-bodies, damaging the membrane.^14 15,16^ We found that Heinz-body formation was highest in HbCC, followed by HbSC and then HbSS, and minimal in HbAA RBC (Figure 1B, Supplemental Figure 2A-D). Notably, HbSS RBC had substantially higher reactive oxygen species (ROS) levels than HbCC and HbSC RBC (Supplemental Figure 2E). The disproportionately higher Heinz-body accumulation in HbC-containing RBC than HbS-containing RBC, despite a higher ROS in HbSS RBC, suggested that HbC is intrinsically more susceptible to oxidative denaturation than HbS. Indeed, HbCC RBC had the highest levels of ferryl (Fe^4^+)-Hb, with HbCC>HbSC>HbSS>HbAA (Figure 1C). This was also associated with the lowest RBC deformability in HbCC<HbSC<HbSS<HbAA (Figure 1D), confirming that the HbC contributes to significant denaturation-induced RBC membrane damage. We next examined Heinz body formation and RBC rheology in HbSC patients to measure hemoglobin denaturation and whether it is modified by hydroxyurea. Hydroxyurea treatment was associated with negligible/modest increases in HbF compared to HbSC patients not on hydroxyurea (4.5% vs. 1.1% HbF and 39% vs. 13% F cells, n=3/group), without changes in hemoglobin, MCV or MCHC (Supplemental Figure 3A-E). There were significantly fewer Heinz-body-positive RBC in hydroxyurea-treated than untreated HbSC patients (Figure 2A, Supplemental Figure 3F), illustrating that hydroxyurea reduces hemoglobin denaturation. Consequently, RBC deformability (determined by the measuring elongation index with increasing shear-stress at normoxia) was higher in hydroxyurea-treated individuals compared to untreated controls (Figure 2B).

**Figure 1.**
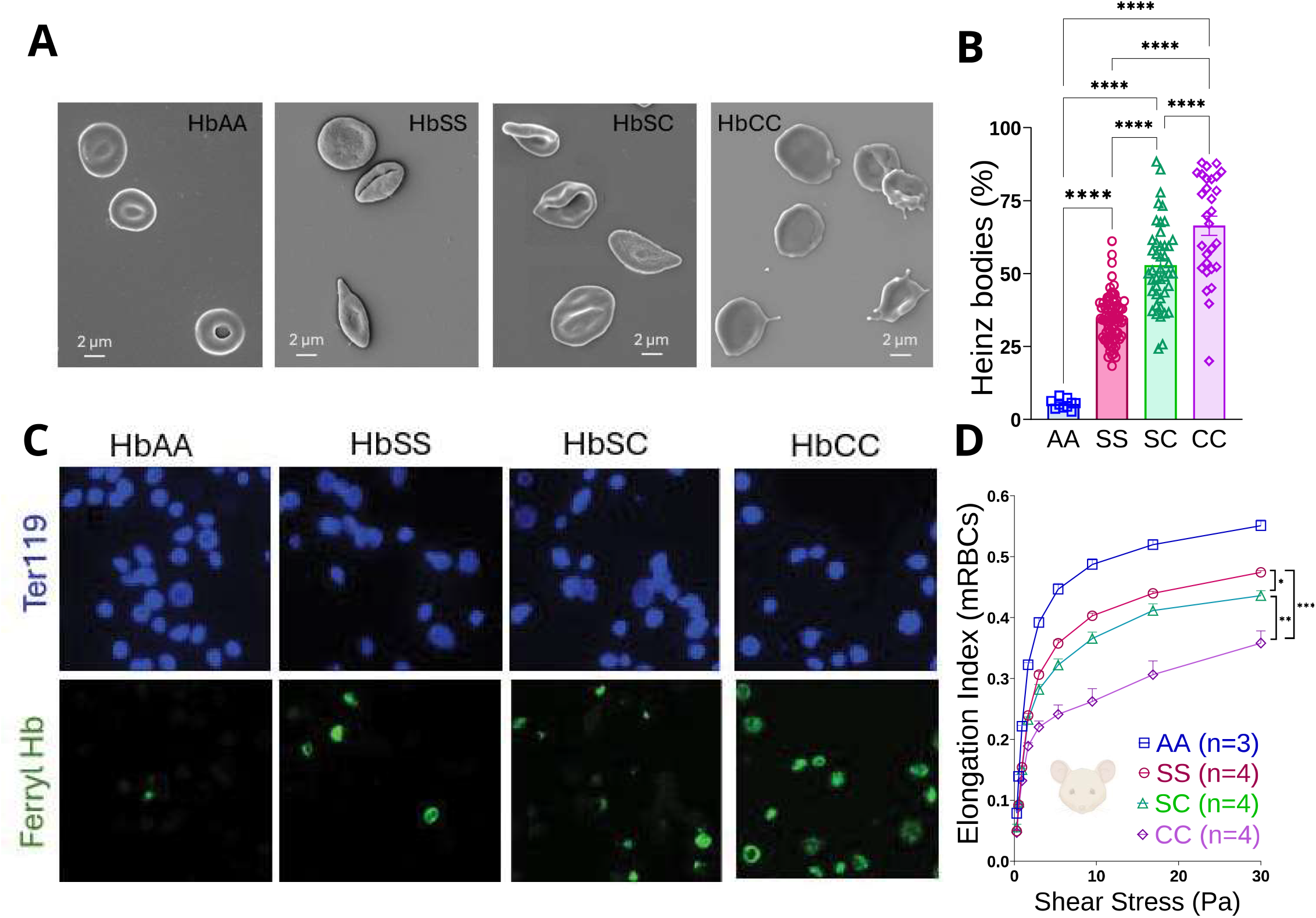
HbC-containing red blood cells exhibit increased hemoglobin denaturation and impaired deformability. **(A)** Representative scanning electron microscopy images of RBCs from HbAA, HbSS, HbSC, and HbCC mice showing distinct morphologic characteristics across genotypes. HbAA RBCs display the typical biconcave discocyte morphology, whereas HbSS, HbSC, and HbCC RBCs show progressively increased membrane irregularities. Specimens were imaged using a Zeiss FE-SEM (Supra 35 VP, Germany) at 2-5 keV with a working distance of 3-5 mm using the lnlens detector. The magnification scale is shown within the micrographs. **(B)** Percentage of Heinz body-positive RBCs across genotypes. HbCC RBCs showed the highest burden of Heinz body formation, followed by HbSC RBCs. Heinz body formation was analyzed in four individual mice for each phenotype. A total of 10-15 high-power fields/blood smear/mouse were quantified (technical replicates), bars represent mean ± SEM. Statistics were performed using ANOVA. ****P<0.00001. **(C)** Immunofluorescence imaging of ferryl Hb in RBCs from HbAA, HbSS, HbSC, and HbCC mice. Anti–ferryl Hb (green) and Ter119 (blue) staining were visualized on an EVOS 7000 (ThermoFisher Scientific) using a 60 × coverslip-corrected objective (Olympus) and analyzed in the EVOS analysis software (ThermoFisher Scientific). Scale bar, 10 µm. **(D)** Shear-stress ektacytometry, which measures RBC deformability, expressed as the elongation index (El), across increasing shear stress levels in HbAA, HbSS, HbSC, and HbCC mice.

**Figure 2.**
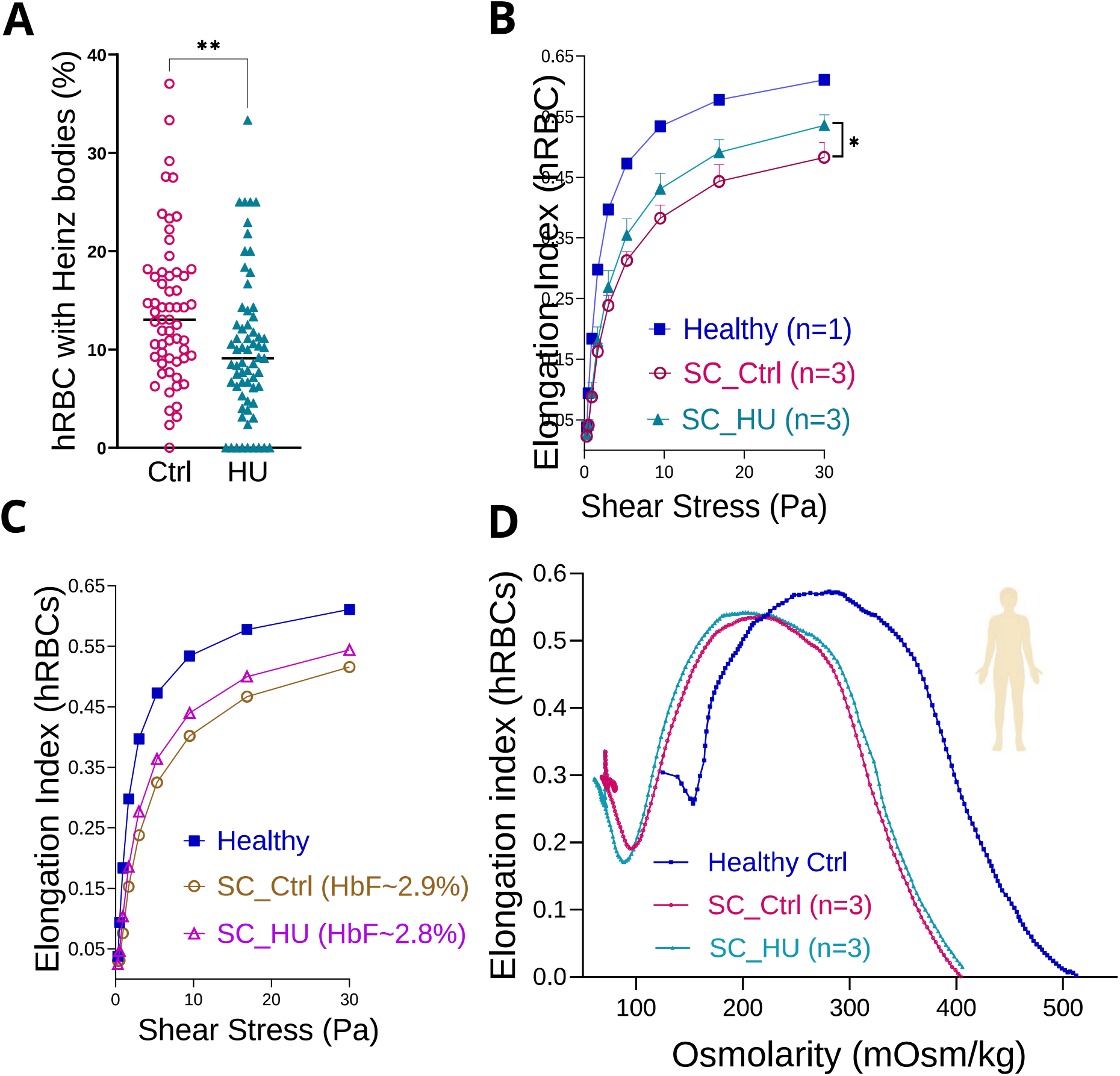
Hydroxyurea reduces oxidative hemoglobin denaturation and improves RBC mechanics in HbSC patients. **(A)** Quantification of Heinz body–positive RBCs in peripheral blood smears from untreated and hydroxyurea-treated HbSC patients. Heinz body formation was measured in 3 HbSC patients receiving hydroxyurea and 3 untreated patients. A total of 20-25 high-power fields/blood smear/patient were quantified (technical replicates). Each point represents an individual measurement; bars indicate mean ± SEM. Statistics were performed using T-Test. * P<0.05, **P<0.01. **(B)** Shear-stress ektacytometry demonstrating improved RBC deformability in hydroxyurea-treated HbSC patients compared with untreated patients (n=3/group). **(C)** Representative shear-stress ektacytometry curves from two human samples with comparable HbF levels (∼2.9%), including one hydroxyurea-treated sample and one untreated control. Elongation index (El) was measured across increasing shear stress levels. **(D)** Osmoscan curves for comparing healthy control RBCs with untreated and hydroxyurea-treated HbSC RBCs under varying osmolar conditions.

This improvement in deformability could result from reduced sickling-induced membrane damage (due to modest HbF increases) or due to reduced hemoglobin denaturation. Representative oxygen-gradient ektacytometry tracings of an untreated HbSC patient with 0.9% HbF and a hydroxyurea-treated HbSC patient with 7% HbF illustrated improved deformability (elongation index [El] maximum [max] and El minimum [min]), with a lower point-of-sickling in the hydroxyurea-treated patient (Supplemental Figure 3G). Comparisons of oxygen-gradient tracings of two HbSC patients with similar (-3%) HbF, one with and the other without hydroxyurea-treatment, showed better RBC deformability on hydroxyurea (Supplemental Figure 3H). A larger sample size is needed to determine the exact effect of HbF versus Heinz-bodies and oxygen-gradient ektacytometry. Hydroxyurea did not affect RBC hydration, as determined by osmotic-gradient ektacytometry (no change in O_hyper_), which was consistent with the lack of change in MCHC (Figure 2D, Supplemental Figure 3E). Together, these data appear to indicate that hydroxyurea reduces oxidative hemoglobin-mediated membrane damage to improve RBC deformability in HbSC disease, and this effect occurs despite modest HbF induction, consistent with our recently published results in HbSC mice^11^ and clinical human data.^9^

To directly test whether reducing oxidative hemoglobin denaturation can improve RBC mechanics independent of HbF, we studied the effects of hydroxyurea and the antioxidant quercetin in HbSC mice (Figure 3A). Quercetin is a natural bioflavonoid that scavenges ROS.^17-19^ In adult humanized HbSC/HbSS mice, the human y-globin gene is transcriptionally switched off prenatally, and HbF protein is no longer detectable after 1-2 weeks of age, nor can it be reactivated with hydroxyurea.^20^ Quercetin or hydroxyurea, alone or in combination, did not significantly change HbF (0% in all groups), F-cells levels, hemoglobin, and reticulocytes, compared to untreated HbSC controls (Supplemental Figure 4A-C). However, in HbSC RBCs, both quercetin and hydroxyurea, alone or in combination, significantly reduced the Heinz-body burden compared with untreated HbSC controls (Figure 3B). Quercetin-treatment of HbCC and HbSS mice also led to reduced Heinz-body burden in their RBC (Supplemental Figure 5A), indicating reduced oxidative hemoglobin denaturation with quercetin. The combination of hydroxyurea and quercetin did not further enhance the effect, suggesting that both agents may act by the same pathway (Figure 3B). We also assessed rheology by oxygen-gradient ektacytometry. The reduction in Heinz-bodies with hydroxyurea or quercetin reduced membrane damage and increased RBC deformability (increased Elmax), but there was no significant improvement in Elmin and point-of-sickling (Figure 3C-I). Notably, neither hydroxyurea nor quercetin changed RBC xerocytosis, analyzed by osmotic gradient ektacytometry (Figure 3J), indicating that Heinz-body-induced membrane damage does not contribute to xerocytosis caused by HbC. Similarly in HbAA, HbSS, and HbCC mice, quercetin improved RBC deformability by shear-stress ektacytometry, but did not improve cellular hydration (Supplemental Figure 5B-F). Collectively, these findings suggest that reducing oxidative hemoglobin denaturation can reduce RBC membrane damage, independent of HbF induction and RBC hydration.

**Figure 3.**
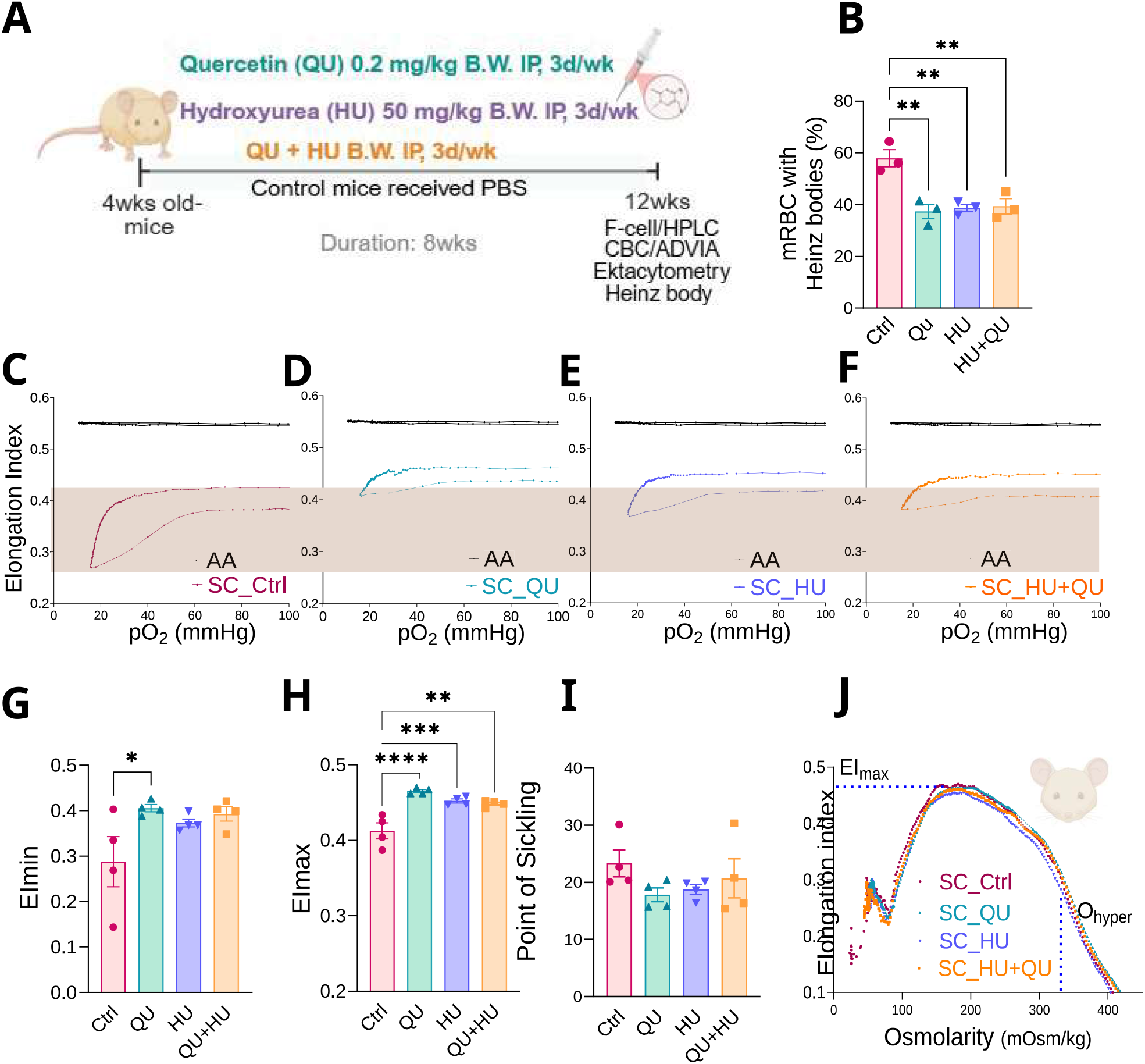
Antioxidant therapy reduces hemoglobin denaturation and improves RBC rheology in HbSC mice. **(A)** Experimental design. Four-week-old humanized HbSC mice were treated for 8 weeks with quercetin (QU; 0.2 mg/kg body weight), hydroxyurea (HU; 50 mg/kg body weight), or the combination (HU+QU), administered intraperitoneally 3 times weekly. Control mice received PBS. Blood was analyzed at 8 weeks for CBC, HPLC, Heinz bodies, and ektacytometry. **(B)** Percentage of Heinz body-positive RBCs in untreated and treated HbSC mice. Each symbol represents an average of 600 RBCs in an individual mouse, and bars represent mean± SEM. **(C-F)** Representative oxygen-gradient ektacytometry curves from untreated HbSC mice **(C)** and HbSC mice treated with QU **(D), HU (E)**, or HU+QU **(F)**, shown in comparison with HbAA reference curves. The shaded area indicates the Elmax-to-Elmin range of control HbSC RBCs to facilitate visual comparison. **(G-1)** Quantification of oxygen-gradient ektacytometry parameters in control and treated HbSC mice, including Elmin **(G)**, Elmax **(H)**, and point of sickling **(I)**. Each symbol represents 1 mouse; bars indicate mean ± SEM. Statistical analysis was performed by one-way ANOVA. * P< 0.05; ** P<0.01; *** P<0.001; **** P<0.0001; ns, not significant. **(J)** Osmotic gradient ektacytometry showing elongation index across osmolarity in control and treated HbSC mice. Neither HU nor QU, alone or in combination, significantly improved the dehydration parameter Ohyper.

HbC is relatively unstable compared with HbA and is susceptible to autoxidation, hemichrome formation, and precipitation on membranes, forming Heinz-bodies,^11,21^ injuring RBC membranes and impairing their deformability.^15,16^ Our data show that HbC is even more susceptible to oxidative damage than HbS, and identify that HbC-driven hemoglobin denaturation is an important determinant of membrane injury and abnormal rheology in HbSC disease.

Our data also show that HbC-induced oxidative membrane damage and xerocytosis are separable components of HbSC pathophysiology, and that HbC-caused xerocytosis occurs by a currently unidentified mechanism. Importantly, our data adds another prong of oxidative hemoglobin denaturation-induced membrane damage alongside HbS polymerization/sickling and cellular dehydration in HbSC disease pathophysiology. While HbS also contributes to membrane injury and dehydration, albeit to a lesser extent than HbC, the combined deleterious effects of HbS and HbC likely explain the substantial morbidity seen in HbSC disease, despite lower HbS content than in HbSS-SCD. It will be interesting to study whether HbF-inducing genetic therapies developed for HbSS-SCD, which primarily target HbS polymerization, also improve oxidative membrane injury.

## Acknowledgments

We thank the Erythrocyte Diagnostic Laboratory at CCHMC, especially Dr. Theodosia Kalfa, Jenifer Korpik, Miya Lu, Ellie Savidge, and Kristina Hanson, for their support with hematological testing. Research Flow Cytometry Core Facility at Cincinnati Children’s Hospital Medical Center (CCHMC) for flow cytometry support, and Abigail Kincaid for HbF quantitation. This work was supported in part by a Center for Clinical and Translational Science and Training Just-in-Time grant, the Research Innovation/Pilot grant, and an American Society of Hematology Bridge Grant to PM and American Society of Gene and Cell Therapy Career Development Award to TS. The graphical abstract and Figure 3A were created with BioRender.

## Author contributions

TS and PM conceived and conceptualized the project. TS, RW, and PM designed the experiments, developed the methodology, interpreted the data, and wrote the manuscript. Data was acquired and analyzed by TS, AT, SK, HK, ZO, KS, JB, MC, and ZZ. Data plotting and micrography were performed by TS and PM. All authors reviewed, commented on, and approved the manuscript.

## Figure Legends

**Supplemental Figure 1.**
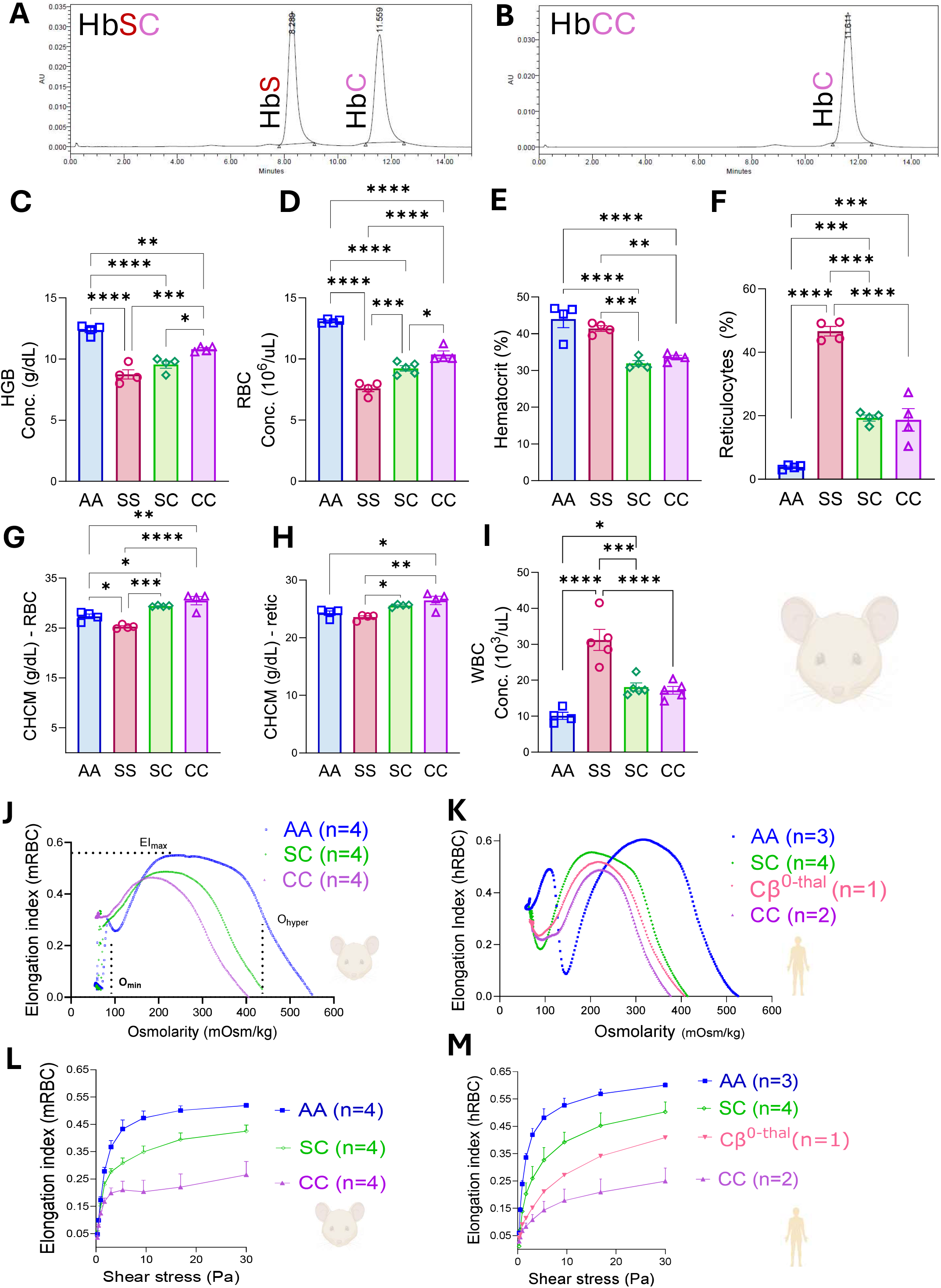
HbCC mice show marked xerocytosis and reduced deformability compared with HbSC, HbSS, and HbAA mice. (**A-B**) Representative cation-exchange HPLC chromatograms confirming the expected human globin composition in HbSC and HbCC mice. HbSC mice show peaks corresponding to both HbS and HbC, whereas HbCC mice show a predominant HbC peak. (**C-I**) Hematologic characterization of HbAA, HbSS, HbSC, and HbCC mice, including hemoglobin concentration (**C**), RBC count (**D**), hematocrit (**E**), reticulocyte percentage (**F**), cellular hemoglobin concentration mean (CHCM) of mature RBCs (**G**) and reticulocytes (**H**), and white blood cell count (**I**). HbCC mice showed prominent erythrocyte abnormalities with increased cellular hemoglobin concentration, consistent with marked RBC dehydration/xerocytosis. Data are presented as mean SEM. Statistical analysis was performed by one-way ANOVA with multiple-comparison testing. (**J-K**) Oxygen-gradient ektacytometry curves comparing HbAA, HbSC, and HbCC RBCs in mice (n=4/phenotypes) (*J*) and HbAA (n=3), HbSC (n=5), HbCβ0-thal (n=1), and HbCC (n=2) RBCs in human (**K**). HbCC RBCs exhibited severely impaired deformability across the oxygen gradient, with greater impairment than HbSC RBCs in mice, similar to humans. (**L-M**) Osmotic gradient ektacytometry showing elongation index across osmolarity in HbAA, HbSC, and HbCC RBCs in mice (n=4/phenotypes) (**L**) and HbAA (n=3), HbSC (n=5), HbCβ0-thal (n=1), and HbCC (n=2) RBCs in humans (**M**). HbCC RBCs showed a left-shifted osmoscan profile and increased Ohyper, consistent with pronounced xerocytosis and reduced membrane deformability.

**Supplemental Figure 2.**
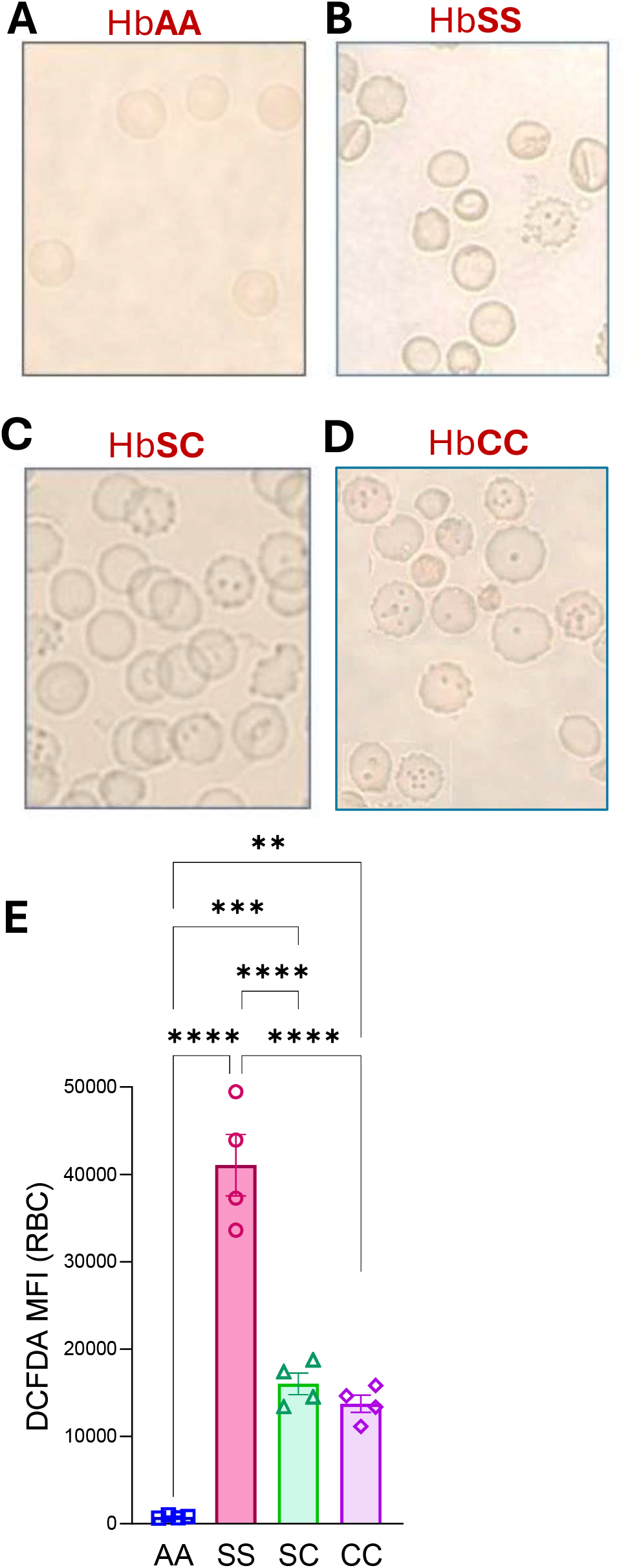
HbC-containing RBCs show marked Heinz body formation despite lower intracellular ROS than HbSS RBCs. (**A-D**) Representative images of Heinz body RBCs from HbAA, HbSS, HbSC, and HbCC mice. (**E**) Quantification of DCFDA mean fluorescence intensity (MFI) in RBCs from HbAA, HbSS, HbSC, and HbCC mice. HbSS RBCs showed the highest intracellular reactive oxygen species (ROS), whereas HbSC and HbCC RBCs had significantly lower DCFDA fluorescence despite marked hemoglobin denaturation. Each symbol represents 1 mouse; bars indicate mean ± SEM. Statistical analysis was performed by one-way ANOVA with multiple-comparison testing. **P < .01; ****P < .0001; ns, not significant.

**Supplemental Figure 3.**
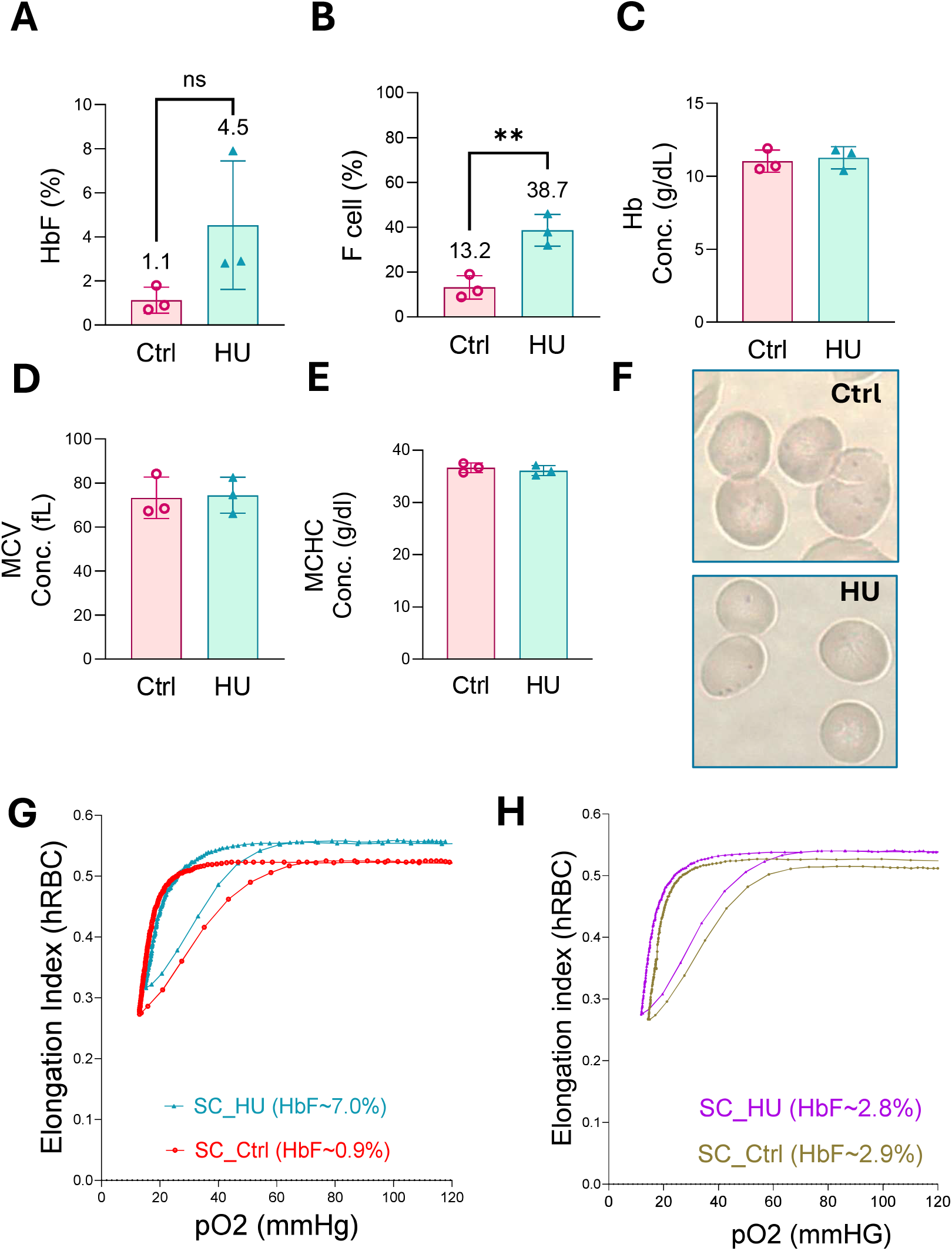
Hematologic parameters in untreated and hydroxyurea-treated patients with HbSC disease. (**A-D**) HbF percentage (**A**), hemoglobin concentration (**B**), % F cell (**C**), mean corpuscular volume (MCV), and (**D**), mean corpuscular hemoglobin concentration (MCHC) (**E**) in untreated (Ctrl) and hydroxyurea-treated (HU) HbSC patients. Each symbol represents 1 patient; horizontal lines indicate mean values. (**F**) Representative image of Heinz bodies in HU and Ctrl (**G**) Representative oxygen-gradient ektacytometry (OxyScan) from the hydroxyurea group with 7% HbF compared to SC untreated with 0.9% HbF, showing improved RBC deformability across decreasing oxygen tensions in hydroxyurea-treated HbSC RBCs. (**H**) Representative oxygen-gradient ektacytometry (OxyScan) from two human samples with comparable HbF levels (∼2.9%), including one hydroxyurea-treated sample and one untreated control, showing that even with no HbF improvement, the Elmax slightly improved.

**Supplemental Figure 4.**
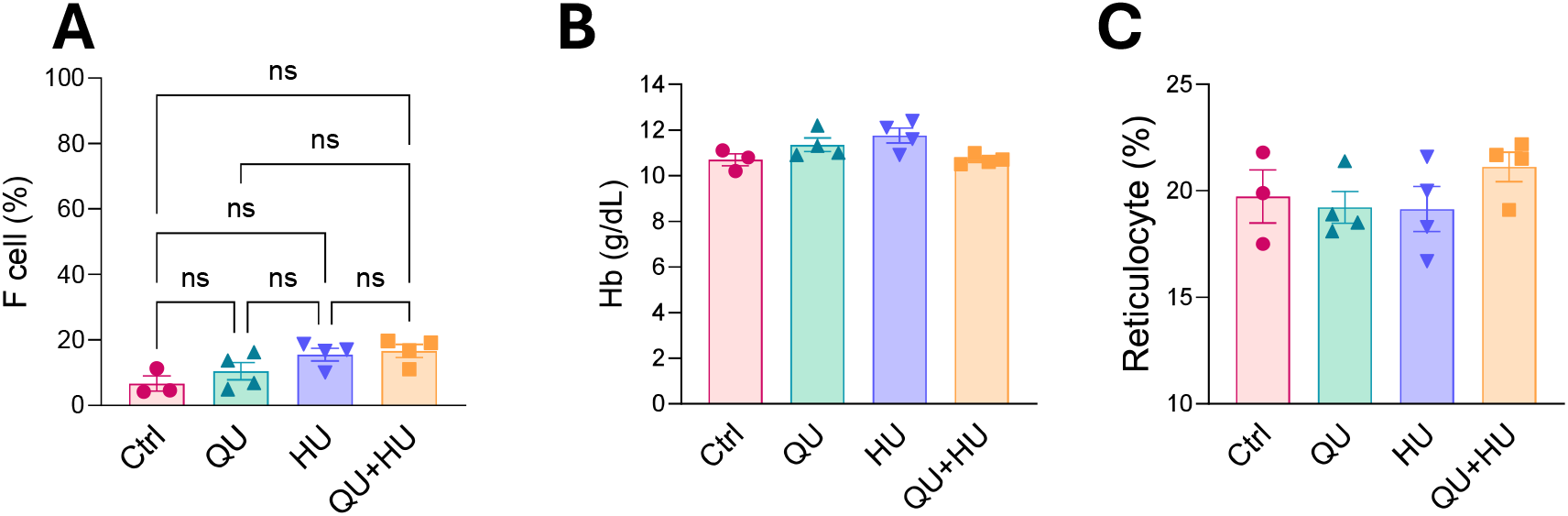
Hematologic parameters and spleen volume are unchanged after hydroxyurea and quercetin treatment in HbSC mice. (**A**) percentage of F cell (**B**) hemoglobin concentration (**C**) Reticulocyte percentage in control HbSC mice and HbSC mice treated for 8 weeks with quercetin (QU), hydroxyurea (HU), or the combination (QU+HU). No significant differences were observed among groups. Each symbol represents 1 mouse; bars indicate mean SEM. Statistical analysis was performed by one-way ANOVA.

**Supplemental Figure 5.**
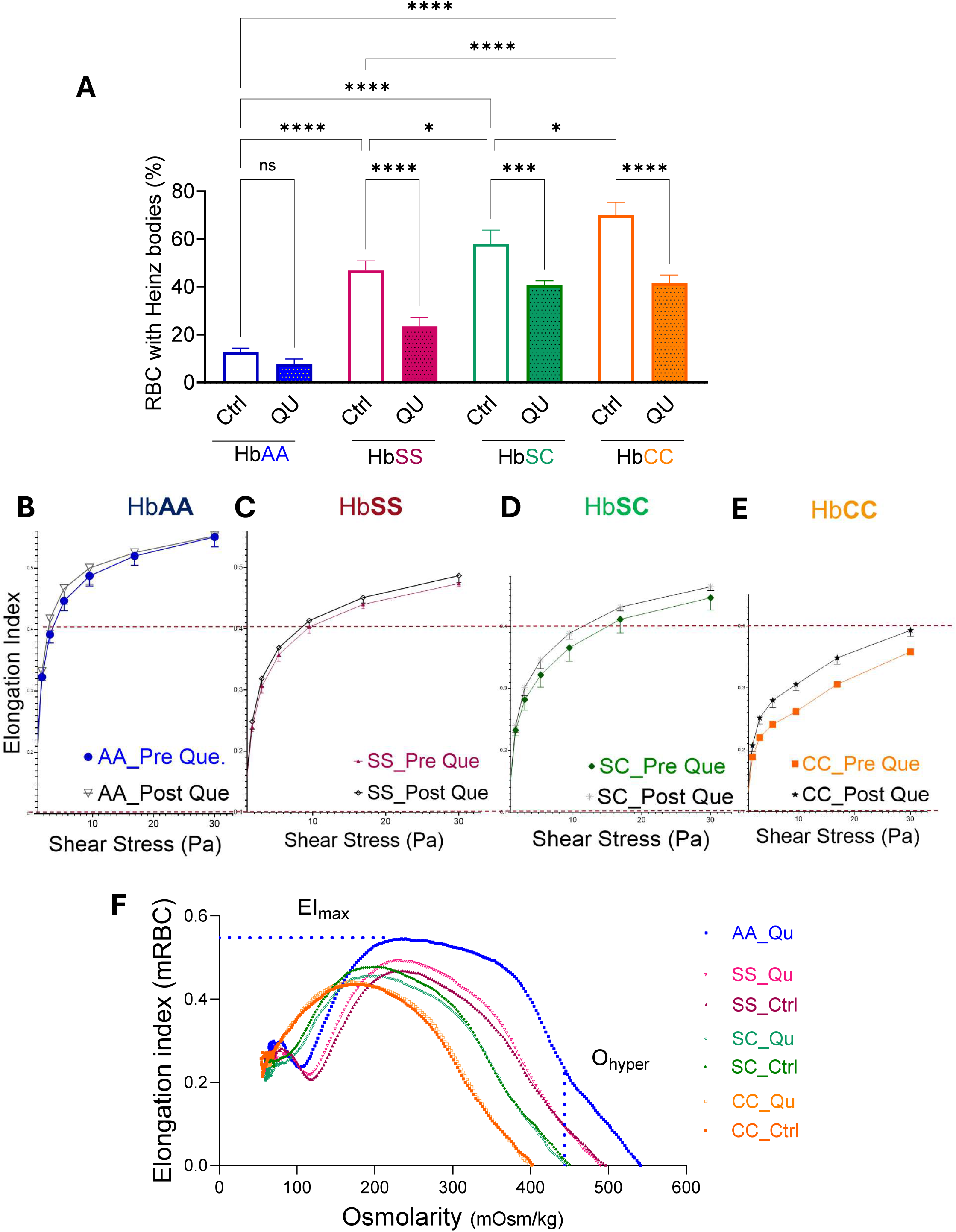
Quercetin reduces Heinz body formation and improves RBC deformability across hemoglobin genotypes without altering hydration-related parameters. (**A**) Quantification of Heinz body–positive RBCs in HbAA, HbSS, HbSC, and HbCC mice treated with quercetin (Que) 0.2 mg/kg, BW, IP, three times a week for 4 weeks or untreated (PBS - Cont.). Quercetin reduced Heinz body formation in HbSS, HbSC, and HbCC RBCs, with the most marked effect in HbC-containing cells. A minimum of 600 RBCs were counted per mouse; 4 mice were analyzed per group. Bars indicate mean SEM. Statistical analysis was performed by one-way ANOVA with multiple-comparison testing. ns, not significant; *P < .05; ***P < .001; ****P < .0001. (B-E) Shear-stress ektacytometry showing RBC deformability across increasing shear stress in HbAA (**B**), HbSS (**C**), HbSC (**D**), and HbCC (**E**) mice before and after quercetin treatment. Deformability is expressed as the elongation index (EI). The dashed horizontal line indicates EI = 0.4 to facilitate comparison across genotypes. (**F**) Osmotic gradient ektacytometry curves from control and quercetin-treated RBCs across genotypes. Quercetin improved deformability in HbSS, HbSC, and HbCC RBCs but did not measurably change the dehydration parameter Ohyper.

